# Spectrum of Opportunistic Disease and Associated Factors among Patients Attending ART Clinic, Nekemte Specialized Hospital, Western Ethiopia

**DOI:** 10.1101/2020.10.06.327668

**Authors:** Meseret Belete Fite, Demeke Jabessa Aga

## Abstract

**Introduction:** Human immunodeficiency virus (HIV), the causal agent for Acquired I Immunodeficiency Syndrome (AIDS) is the world’s greatest severe public health and development contest. Since the beginning of the epidemic, 38 million of people are living with HIV/AIDS and 1.7 million people newly infected with HIV. Increased availability and accessibility of ART has essentially improved the survival rate, through lowering the incidence of OIs among peoples living with HIV/AIDS. Risk of developing Opportunistic infections in HIV patients depend on experience to potential pathogens, virulence of pathogens, degree of host immunity and the use of antimicrobial prophylaxis. In Ethiopia, however remarkable decline of new infection (81%) for decades, since 2008 HIV incidence rate began to rise by 10% and number of new infection diagnosed each year increased by 36% among all ages and doubled among adult. There is a limited studies describing the spectrum of opportunistic infection and associated factors in the study settings. Therefore, this study was aimed to determine the spectrum of Opportunistic infections in the study area.

**Methods:** A Facility based retrospective cross-sectional study was employed from 2015-2019 G.C. The sample size was computed using single population proportion formula. Accordingly, four hundred ninety seven (497) medical records of study participants were reviewed. Simple random sampling technique was used to select the participants included in this study. Data were extracted from the ART follow up data-base and medical records of the patients by using a standardized check list, which was adapted from Federal ministry Of Health **HIV ART**. The contents of check list include: Socio-demographic characteristics and clinical information’s. Data had entered Epi data version and analyzed using SPSS version 5.3.1 and analyzed using SPSS version 20. Bivariate analysis with p-value <0.2 was done to see the association between outcome variable & independent variables. Variables with p < 0.2 in bivariate analysis were entered for multiple logistic regressions. At 95% confidence interval, explanatory variables with P <0.05 in multiple logistic regression analysis were considered as significantly association.

**Result:** The study found that, an overall prevalence of OIs was 62%. The finding of our study documented, from deferent HIV related OIs among patients on ART follow up at Nekemte Specialized Hospital ART clinic, the common types of OIs were; Pulmonary Tuberculosis (15.7%), Oral candidiasis (14.3%), Herpes Zoster (11.3%), Cryptococcus meningitides (5.9%), upper respiratory infection (5.8%, Persistent diarrhea (5.2%), and Extra pulmonary tuberculosis (3.8%). The occurrence of OIs on adult PLHIV patients who were with baseline WHO stage of I were 53% lower as compared to those who were with advanced baseline WHO stage of II and more {AOR**: 0**.**468**, 95 % CI (**0**.**305-0**.**716). Moreover**, Participants of Urban residents were 1.6 times more likely to develop OIs than those rural residents. Baseline WHO clinical staging and residence were identified as independent predictors of OIs among adult HIV infected patients.

**Conclusion:** An overall prevalence of OIs was 62%. The prevalence of OIs is still high namely Pulmonary Tuberculosis, Oral candidiasis and Herpes Zoster are leading OIs among adult HIV infected patients. Baseline WHO clinical staging and residence were identified as independent predictors of OIs among adult HIV infected patients

**Recommendations:** Having skilled health professionals, early diagnosis of OIs among HIV infected patients and having equipped laboratory diagnostic setup are mandatory to be able to deal with specific diagnosis and management of OIs. Further study is recommended to determine the relationship between residence and developing OIs among HIV patients on ART follow

## Introduction

Human immunodeficiency virus (HIV), the causal agent for Acquired I Immunodeficiency Syndrome (AIDS) Is the world’s greatest severe public health and development contest (1). As per global joint estimate of World Health Organization and United Nation Program on HIV/AIDS 2019, since the beginning of the epidemic, 76 million people have been infected with the HIV virus and about 33 million people have died of HIV/AIDS, 38 million of people are living with HIV/AIDS; 1.7 million people newly infected with HIV in 2019 and 690,000 peoples were died with HIV related diseases(2). This epidemic remains Global public health challenge for 21 century(3), in the absence of vaccine and curative therapy. Opportunistic infections (OIs) are defined as infections that occur more often or are more severe in people with weakened immune systems than in people with healthy immune systems because of immune suppression in HIV infected person, and they are major clinical manifestation of HIV patents (4). Increased availability and accessibility of ART has essentially improved the survival rate, through lowering the incidence of Opportunistic infections among peoples living with HIV/AIDS(5). In Ethiopia, however remarkable decline of new infection (81%) for decades, since 2008 HIV incidence rate began to rise by 10% and number of new infection diagnosed each year increased by 36% among all ages and doubled among adult (6)

Risk of developing Opportunistic infections in HIV patients depend on experience to potential pathogens, virulence of pathogens, degree of host immunity and the use of antimicrobial prophylaxis(7). Opportunistic infections (OIs) associated with HIV remain the single main cause of ill-health and death among HIV/AIDS patients in resource poor settings, therefore HIV positive patients in resource poor settings continue to suffer from(8). Evidence of studies conducted in different parts of Ethiopia had documented that, the incidence of OIs after HAART was increased in especially among hospitalized HIV-patients (9,10) and, reported the prevalence of OIs among HIV infected patients in the range of 33.3% in Addisa Abeba to 88% in Dawro Zone Hospital 11-16). Tuberculosis (TB), oral candidacies, Pneumocystis Carini Pneumonia (PCP), bacterial pneumonia, Herpes Zoster, Cryptococcus meningitides, Persistent diarrhea, kaposi’s sarcoma and lymphoma are the most reported common opportunistic infections in HIV patients in Ethiopia (15,17,18), but major differences in spectrum of OIs observed across the country.

Various literatures suggested that, history of opportunistic infections, Hgb levels, WHO clinical staging, CD4 counts, taking past OI prophylaxis, ART drug adherence, monthly income and occupation were identified as predictors of developing OIs among HIV infected patients(14-16,19,20). However, methodological difference and variation of duration of HART were observed across these studies. Timely specific diagnosis and consequent treatment to combat these infections is not only the concern for industrialized countries, but also in the same way important to developing countries like Ethiopia. In Ethiopia, early diagnosis of OIs in HIV infected patients is not an easy task, as there is scarcity of laboratory diagnostic setup to be able to deal with specific diagnosis of opportunistic infections. Therefore, the present study was planned in order to assess the prevalence, spectrum and predictors of OIs among adult HIV infected patients in this setup.

## METHODS

### Study setting

The study was conducted in Nekemte Specialized Hospital ART clinic. This is located in Nekemte town, East Wollega Zone at the distance of 328 km to the west of the capital city of Ethiopia., Addis Ababa. Nekemte Specialized Hospital is the only public specialized hospital in, Western Oromia region. The hospital provides preventive, curative, and rehabilitative services for populations of more than 3.5 million of Western Zones of Oromia region. Also three governmental and one nongovernmental health centers are serving the population of the town. ART service for Nekemte Specialized Hospital was initiated in 1998 G.C and has one clinic. A total of 7200 adult patients were enrolled in ART starting from Jan, 2015-Jan, 2019 G.C

### Study design and period

Facility based retrospective cross-sectional study was conducted from Jan, 2015-Jan, 2019 G.C

### Source population and study population

The source population was all adult patients who were on ART follow up in Nekemte specialized hospital, ART clinic. The study population was all adult patients who were selected by systematic sampling method from source population and participated in the study during study period.

### Inclusion and Exclusion Criteria

Clinical records of all patients who were on ART follow up and 18 years and above during study period were included. However, Clinical records that do not have complete information relevant for the study were excluded from the study.

### Sample Size Determination

The sample size was computed using single population proportion formula shown below. Prevalence of opportunistic infection among adult patients on ART follow up (P= 48%) in Eastern Ethiopia was taken to determine sample size (13). Margin of error (5%) and critical value at 95 % confidence level was used. Ten percent (30 %) contingency for incomplete data was considered. Finally, 498 adult patients on ART follow up were included in this study.

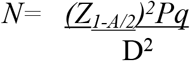

### Sampling Procedure and technique

Nekemte Specialized Hospital was purposefully selected for the study. The hospital had one ART clinic. A total of 7200 patients were enrolled on HIV starting from Jan, 2016-Jan, 2019 G.C. Then, using simple random sampling technique; a computer-based random number generating using the patient’s registry as sampling frame, a total number of 498 patients were selected to be included in the study. Data of 2015-2019 was considered and any incidence of OIS and co-morbidities was collected both from patient chart and history sheet.

### Operational Definition

**Development of OIS**:-infections caused secondary to HIV infection.

**Adherence level**: -Is the degree to which patients accept treatment process

**Adherence**:-is defined as good if adherence is > 95% (< 2 doses of 30 doses or < 3 dose of 60 dose is missed); poor if adherence is between 85 and 94% (3–5 doses of 30 doses or 3–9 dose of 60 dose is missed). Adherence was measured by pills count.

### Data collection tools and techniques

Data were extracted from the ART follow up data-base and medical records of the patients by using a standardized check list, which was adapted from Federal ministry Of Health **HIV ART**, follow up guideline {21}. The contents of check list include: Socio-demographic characteristics (age, sex, marital status, and Residence, ethnicity, religion, and employment & Educational status) and Clinical information’s (CD4 level, WHO stage, Adherence, weight, co-morbidities). Data extraction was performed by trained health professional that had experience of ART clinic service. Total of 3 individual two BSc nurse and 1 Health officer were involved for data extraction. Data was retrieved from the hospital’s ART registry document which is a standard format for sending comprised data to health monitoring information system.

### Data Quality Assurance

Two day training was given for each data collectors and supervisors. The training was also emphasized on the importance of the privacy or confidentiality of patient. Data collectors were supervised by ART clinician. Regular supervision was given during data collection. Collected data was checked by supervisors before sent to the data entree on daily basis. Data collection material was pretested using similar set up at Nekemte health center. To ensure the quality of data, selected individual’s medical record was evaluated for completeness and all the needed information were checked. To avoid bias, Patient clinical records with redundant, incomplete or missing information were omitted.

### Data Processing and Analysis

Data was first checked manually for its completeness and consistency by supervisors & investigators during the time of data collection and rechecked again before data entry. The prevalence of OIs was determined as the proportion of HIV/AIDS patients on ART who developed one or more OIs. Descriptive statistics was done to summarize the data. In this study adult patients who were on ART follow up and developed either one or more of OIs (pulmonary tuberculosis, Oral candidiasis, Herpes Zoster, Cryptococcus meningitides and upper respiratory infection, persistent diarrhea, extra pulmonary tuberculosis, minor mucocutaneous infection, recurrent bacterial infection, Esophageal candidiasis, lymphoma and Toxoplasmosis of Central nerve system) were considered as hiving Opportunistic infection. Data had entered Epi data version 5.3.1 and analyzed using SPSS version 20. Frequencies, cross tabulation and percentages, were calculated for all the categorical variables. Then bivariate analysis with p-value <0.2 was done to see the association between outcome variable & independent variable.. Binary logistic regression analysis was conducted. Variables with p < 0.2 in bivariate analysis were entered for multiple logistic regressions. At 95% confidence interval, explanatory variables with P <0.05 in multiple logistic regression analysis were considered as significantly association.

### Ethics Considerations

Ethical clearance was obtained from the Institutional Health Research Ethics Review Committee of Oromia Regional Health Bureau, Ethiopia. Then, official letter was written to Nekemte Specialized Hospital for permission. Nekemte specialized hospital management and staff of ART clinic were requested for permission of entry using an official letter and the hospital granted permission for data collection. Patient’s’ name were not collected. All data extracted from the patient’s registry were kept strictly confidential. Since secondary data was used, consent was waived by hospital management and staff of ART clinic. The HIV infected, the result of the study was communicated to respective any beneficiary bodies

## Result

### Socio-Demographic Characteristics

A total of four hundred ninety seven study adult HIV/AIDS patient’s ART records were looked over in this study. The mean age of the respondents was 33 (**SD ± 8)** years and ranging from 25-41 years. Nearly about half 247(49.7%) of the study participants were in an age interval of 26-37 years. From total respondents majority, 312(62.8%) of them were females and, 308(62%) were living in urban area. Regarding to marital status, more than half of the study participants, 311(62.6%) were married and 186(37.4%) of them were single. Among the participants, 242(48.7%) were orthodox religion followers and, 178(35.8%) had attended primary education. With regard to the educational status, most the participants, 293(58.8%) were unemployed (**Table 1**).

**Table: 1.** Socio-demographic characteristics of the study participants (N=497) at Nekemte Specialized Hospital ART clinic, Nekemte, Western Ethiopia, 2020.

| Characteristics | Frequency |  |
| --- | --- | --- |
|  | Number (N) | Percentage (%) |
| **Age category** |  |  |
| 18-25 | 102 | 20.5 |
| 26-35 | 247 | 49.6 |
| 36-45 | 110 | 22.4 |
| 46-55 | 38 | 7.5 |
| **Sex** |  |  |
| Male | 185 | 37.2 |
| Female | 312 | 62.8 |
| **Residence** |  |  |
| Urban | 308 | 62.0 |
| Rural | 189 | 38.0 |
| **Marital status** |  |  |
| Married | 311 | 62.6 |
| Single | 186 | 37.4 |
| **Religion** |  |  |
| Orthodox | 242 | 48.7 |
| Protestant | 205 | 41.3 |
| Muslim | 50 | 10.0 |
| **Educational status** |  |  |
| Primary | 178 | 35.8 |
| Secondary | 141 | 28.4 |
| Tertiary | 84 | 16.9 |
| No formal education | 94 | 18.9 |
| **Occupation** |  |  |
| Employed | 144 | 29 |
| Unemployed | 293 | 58.8 |
| Others | 63 | 12.2 |

### Clinical Characteristics of PLWH

As noted in Table 2, majority of the study participants had good adherence 476(95.8%). and 303(61%) of the patients had developed opportunistic infection. Regarding to current status of the patients majority of them, 420 (85%) were in WHO stage.

**Table 2.**
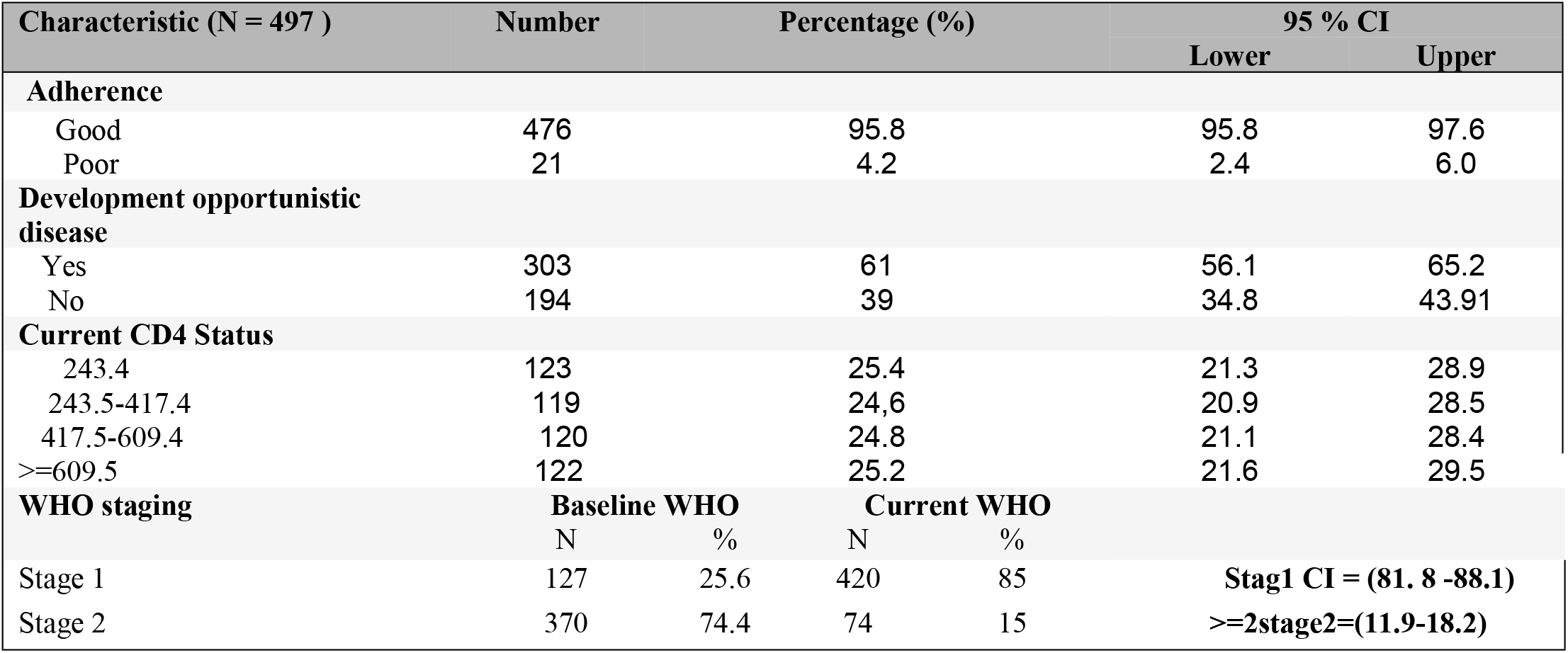
Clinical characteristics of PLWH among clients attending ART clinic (N=497) at Nekemte Specialized Hospital, Nekemte, Western Ethiopia, 2020.

### Prevalence and Spectrum of Opportunistic Disease among PLHIV on ART

Among 497 study participants, 310 had diagnoses of opportunistic infections (OIs), yielding an over prevalence of 62% (310/497). From these, 215(79%) of the HIV infected patients had only one opportunistic infections. Whereas, 45(17%) and 11(4%) of patients had two and three opportunistic infections respectively (Table 3). The most, common type of opportunistic infections among VIV infected patient’s attending ART clinic in the current study were, pulmonary tuberculosis 78(15.7%), followed by Oral candidiasis 71(14.3%), Herpes Zoster 56(11.3%), Cryptococcus meningitides was 29(5.9%), upper respiratory infection 28(5.8%, Persistent diarrhea 27(5.2%), and Extra pulmonary tuberculosis 17(3.8%)(Table 4).

**Table 3:** Number Opportunistic infection (OIs) among adult PLWH at Nekemte Specialized Hospital ART clinic, Nekemte, Western Ethiopia, 2020.

| S/N | Opportunistic infection (OIs) | Frequency (N) | Percentage (%) |
| --- | --- | --- | --- |
| 1 | One Opportunistic infection (OIs) | 215 | 79 |
| 2 | Two Opportunistic infection (OIs) | 45 | 17 |
| 3 | Three Opportunistic infection (OIs) | 11 | 4 |
| 4 | **Total** | **310** | **100** |

**Table 4:** Prevalence of Opportunistic infection (OIs) among adult PLWH at Nekemte Specialized Hospital ART clinic, Nekemte, Western Ethiopia, 2020.

| S/N | Opportunistic infection (OIs) | Frequency (N) | Percentage (%) |
| --- | --- | --- | --- |
| 1 | pulmonary tuberculosis | 78 | 15.7 |
| 2 | Oral candidiasis | 71 | 14.3 |
| 3 | Herpes Zoster | 57 | 11.3 |
| 4 | Cryptococcus meningitides | 30 | 5.9 |
| 5 | Upper respiratory infection | 29 | 5.8 |
| 6 | Persistent diarrhea | 26 | 5.2 |
| 7 | Extra pulmonary tuberculosis | 19 | 3.8 |

**Table 5:** Multiple logistic regression analysis for selected variables with occurrence of opportunistic Infections among adult PLWH at Nekemte Specialized Hospital ART clinic, Nekemte, Western Ethiopia, 2020.

| Characteristic<br>N = 497 | Development of OIS |  | Crude OR with( 95 % CI) | Adjusted OR (95% CI ) |
| --- | --- | --- | --- | --- |
|  | Present | Absent |  |  |
| Adherence |  |  |  |  |
| Good | 288 | 188 | 0.613(0.234 – 1.607) | 0.555(0.200 -1.55) |
| Poor | 15 | 6 | 1 | 1 |
| Current CD4 status |  |  |  |  |
| <243.4 | 78 | 47 | 1.159(0.697-1.929) | 1.235( 0.712 - 2.140) |
| 243.5-417.4 | 77 | 46 | 1.169(0.701-1.950) | 1.332( 0.774 – 2.294 ) |
| 417.4 -609.4 | 74 | 50 | 1.034 (0.623 -1.716 | 1.104 ( 0.651 -1.874 ) |
| >=609.5 | 73 | 51 | 1 | 1 |
| Religion |  |  |  |  |
| Orthodox | 151 | 91 | 0.56(0.146-2.096) | 0.440(.110 - 1.764 ) |
| Protestant | 118 | 87 | 0.452(0.119-1.719) | 0.572(.142 - 2.297 ) |
| Muslim | 25 | 13 | 0.641(0.148 – 2.784 ) | 0.608(.132 -2.798 ) |
| Others | 9 | 3 | 1 | 1 |
| **Residence** |  |  |  |  |
| Urban | **198** | **110** | **1.44 (0.995-2.084 )** | **1.623( 1.097 - 2.399 )\*\*** |
| Rural | **105** | **84** | 1 | 1 |
| **Current WHO stage** |  |  |  |  |
| **WHO Stage I** | 160 | 160 | 1.308(0.794 -2.154) | 1.313(.770 - 2.237) |
| **Stage II and more** | 33 | 33 | 1 | 1 |
| Base line WHO stage |  |  |  |  |
| Stage I | 57 | 62 | 0.497(0.327 –0.755 ) | **0.468(0.305-0.716) \*\*** |
| Stage II and more | 237 | 128 | 1 | **1** |

Reference category , \*\* associated variables ,

### Factors Associated With the Development Of Opportunistic

Multiple binary logistic regression analysis was run, considering those variables p value ≤ 0.25. Factors associated with occurrence of OIs among adult HIV infected patients who were on ART were assessed in the current study. Accordingly, baseline WHO clinical staging and residence were identified as independent predictors of OIs among adult HIV infected patients. The occurrence of opportunistic infection on adult PLHIV patients who were with baseline WHO stage of I were 53% lower as compared to those who were with advanced baseline WHO stage of II and more {AOR**: 0**.**468**, 95 % CI (**0**.**305-0**.**716)**. Moreover, Participants of Urban residents were 1.6 times more likely to develop opportunistic infections than those rural residents.

## Discussion

The current study assessed the prevalence and predictors of opportunistic infection among adult patients who were on ART follow up in Nekemte specialized hospital, ART clinic. The study found that, an overall prevalence of opportunistic infections was 62%. This finding is comparably higher than the prevalence of HIV related opportunistic infections reported in different studies conducted in Addis Ababa (33.3%), North West Ethiopia (33.3%), Eastern Ethiopia (48%), Southern Zone Tigray (55.3%) and Eastern Zone of Tigray (52%)(12-16), Nevertheless comparably lower than of the studies in conducted in Dawro Zone Hospital (88%) and Kenya (78.8%) (11,22). The reason for variation across studies might be due the deference in duration of follow up, difference in CD4 level and degree of host immunity of study subjects. The current study followed study participants for five years.

Evidences of Studies conducted indifferent parts of Ethiopia noted that, opportunistic infections are common among HIV infected people. The finding of our study documented, from deferent HIV related OIs among patients on ART follow up at Nekemte Specialized Hospital ART clinic, the common types of OIs were; Pulmonary Tuberculosis (15.7%), Oral candidiasis (14.3%), Herpes Zoster (11.3%), Cryptococcus meningitides (5.9%), upper respiratory infection (5.8%, Persistent diarrhea (5.2%), and Extra pulmonary tuberculosis (3.8%). Pulmonary Tuberculosis was the most common leading OI among HIV-infected people in this study area. Similarly, study conducted in Dawro Zone hospital, Pulmonary tuberculosis (18%) was observed as most common OI among patients on ART follow up during the study period (11). However the result is comparably lower than finding of study conducted in Southern Zone Tigray (16). In current study Oral candidiasis was observed as the second leading OI; in contrary it was documented as the first common type of OI in study conducted in Debra Marko’s Referral Hospital (11.8%), Southern Zone Tigray (11%), Uganda (34.6%) and India (49%) (12,16,23,24) and, third in Dawro Zone (15.6%) (11). The reason for this variation might possibly be explained by deference in quality of laboratory diagnosis in most part of the country, difference in Geographical area of study participants and methodological deference in selecting of study participants. Herpes Zoster was acknowledged as third common type of OI in this study setup. The prevalence of Herpes Zoster in current study is comparably consistent with the finding of study conducted in Eastern Ethiopia (11.2%)(13) and Southern Zone Tigray(10.8%,) (16), in contrary study of Eastern Ethiopia and Southern Zone Tigray had observed as the second common type of OIs.

This study assessed the level of ART adherence among adult HIV patients who were on ART in Nekemte Specialized Hospital. The level of adherence identified here was 95%, which is consistent with the suggested level of ART treatment adherence. In current suggestion, at least 95% of ART adherence level is needed to overwhelm viral replication, show clinical improvement and better CD4 count (25). Despite, the level of adherence is in agreement with recommended level; the result of current level of adherence was comparably higher than the documented finding of study conducted in Eastern Ethiopia (85%) (20) and Goba hospital (90.8%) (26) and lower than finding of study conducted in Nigeria(98.7%) (17). The reason for the variation could be due to duration of HART. Here in this study, the study participants have been on ART for longer duration, and commonly those who were taking the drug/s for longer period attain skills in what way to compact with difficulties hampering them not to adhere. Further, the context of Socio-Demographic variations might explain the observed difference.

Factors associated with occurrence of OIs among adult HIV patients who were on ART were as well assessed. Concerning factors associated with occurrence of OIs, baseline WHO clinical staging and residence were identified as independent predictors of OIs according to the result of current study. The occurrence of opportunistic infection on adult PLHIV patients who were with baseline WHO stage of I were 53% lower as compared to those who were with advanced baseline WHO stage of II and more. Thus participants with advanced baseline WHO stage of II were 47% more likely risky to develop OIs as compared to those with WHO stage of I. The finding is in agreement with other four studies conducted in deferent parts of Ethiopia (Eastern Ethiopia, Gonder, Addis Ababa and Debre Marcos (12-16,18). Study conducted in Eastern Ethiopia reported that, participants with advanced WHO stage of III were four times more likely to develop OIs than those with WHO stage of I. similarly, Study conducted in Gonder documented that, WHO clinical stage of III HIV infected patients were nine times more likely to develop OIs than those with WHO clinical stage of I. The finding of Study conducted in Debre Marcos noted that, WHO stage of III HIV infected were five times more likely to develop OIs than those with WHO stage of I. likewise, Study conducted in Addis Abeba reported that, HIV infected patients with WHO clinical stage of III were two times more likely to develop OIs than those with WHO clinical stage of I. Comparable conclusion were appreciated in studies conducted in different countries (19,22,23,27

Moreover, an Odd of the occurrence of opportunistic infection on adult PLHIV patients who were urban residents was 1.6 times higher as compared to those who were rural residents. Participants of Urban residents were 1.6 times more likely to develop opportunistic infections than those rural residents. However, studies conducted in deferent in different countries found as no association of residence with developing opportunistic infection (12-13,19,27). Thus limited evidence was generated on association of residence and developing OIs among HIV infected people. Therefore, further study is recommended to determine the relationship between residence and developing opportunistic infections among HIV patients on ART follow up.

## Limitations

Retrospective nature of the study makes some variables less explanatory and cause-effect of relationships cannot be measured. Finally, since we used secondary data some variables were not documented well and many OIs were supposed diagnosis may be stated as likely limitation of current study.

## Conclusion

An overall prevalence of opportunistic infections was 62%. The prevalence of opportunistic infections is still high namely Pulmonary Tuberculosis, Oral candidiasis and Herpes Zoster are leading OIs among adult HIV infected patients. Baseline WHO clinical staging and residence were identified as independent predictors of OIs among adult HIV infected patients

## Recommendation

Having skilled health professional, early diagnosis of OIs among HIV infected patients and having equipped laboratory diagnostic setup are mandatory to be able to deal with specific diagnosis and management of opportunistic infections. Further study is recommended to determine the relationship between residence and developing opportunistic infections among HIV patients on ART follow.

## Abbreviations

HIV: Human immunodeflciency virus
AIDS: Acquired Immune Deflciency Syndrome
ART: Highly Active Antiretroviral Therapy
OIs: Opportunistic Infections
PLWHIV: People Living with HIV
WHO: World Health Organization
USAIDS: United Nation Program on HIV/AIDS

## Declarations

### Consent for publication

#### Availability of data and materials

Data will be available upon requesting of the corresponding author

#### Competing interests

The authors declare no conflicts of interest in this work.

#### Funding

There is no received grant from any fund agency and covered by authors.

#### Authors’ contributions

All authors made substantial contribution from the conception, and design, analysis and interpretation of the data, participated in the drafting and revising of the article critically, and gave final approval of the version to be published, and agree to be accountable for all aspects of the work

## Acknowledgment

Our special gratitude goes Nwkemte Specialized hospital administrator for granted permission for data collection.

